# Transcriptome-wide Differential Expression and Splicing in Inflammatory Bowel Disease Patients

**DOI:** 10.64898/2026.10.24.759415

**Authors:** Shuying Han, Emma Yu-Chen Huang, Sophie Silverstein, Caroline Beasley, Muneera R. Kapadia, Gwen Lau, Grace Lian, Reza Rahbar, Timothy S. Sadiq, Matthew R. Schaner, Jonathan Stem, David Weaver, Benjamin Huan, Ariana LaCorte, Shehzad Z. Sheikh, Terrence S. Furey

**Author notes:** Corresponding authors: Terrence S. Furey, Shehzad Z. Sheikh (shehzad).

## Abstract

The inflammatory bowel diseases (IBD), comprising mainly Crohn’s disease (CD) and ulcerative colitis (UC), are characterized by chronic intestinal inflammation driven in part by complex genetic and gene regulatory mechanisms. Although transcriptomic studies have defined broad expression changes in IBD, the contribution of alternative splicing to disease-associated molecular variation remains less clear. We therefore integrated differential expression (DE) and differential splicing (DS) analyses in macroscopically uninvolved colonic mucosal biopsies from patients with either CD or UC along with non-IBD controls to define gene expression and intron-usage changes across disease subtypes. We identified 491 DS and 316 DE genes in CD vs. non-IBD, and 1,298 DS and 1,107 DE genes in UC vs. non-IBD. DS and DE genes were nearly mutually exclusive, with no overlapping genes in CD and 16 in UC. Consistent with this result, pathway analysis showed that differentially expressed and differentially spliced genes were associated with related but distinct biological programs. DE genes were involved in chromatin organization, transcriptional regulation, and metabolic pathways for glycans and drugs, whereas DS genes were enriched for splicing, cellular signaling, and lipid metabolism. We found substantial similarity in enriched pathways across CD and UC with some notable exceptions, including increased enrichment of DS and DE genes involved in extracellular vesicles and exosomes in CD and enrichment of DS genes in ER-to-Golgi trafficking and stress granules in UC. Direct comparisons of CD and UC patients additionally show differential splicing usage in genes involved in cell adhesion pathways. These findings identify alternative splicing as a major contributor to transcriptomic changes in IBD that is largely distinct from changes in expression level and provide a broader view of molecular contributors to IBD.

## Introduction

The inflammatory bowel diseases (IBD), primarily consisting of Crohn’s disease (CD) and ulcerative colitis (UC), are characterized by epithelial barrier dysfunction, microbial dysbiosis, and dysregulated immune activation, which together drive persistent intestinal inflammation and tissue injury (Falloon and Lazarev, 2019; Neurath, 2019). Epidemiological studies highlight the important contribution of environmental exposures to disease onset and progression, while genetic studies demonstrate that inherited factors play a central role in susceptibility to IBD (Z. Liu et al., 2023; Ananthakrishnan et al., 2017). Although CD and UC share many clinical manifestations, they differ in their anatomical distribution and depth of inflammation. CD can affect any segment of the gastrointestinal tract in a discontinuous pattern and often involves transmural inflammation, while UC is restricted to the colonic mucosa and progresses continuously from the rectum proximally (Younis, Zarif, and Mahfouz, 2020). These pathological differences suggest that the two diseases may arise from partially distinct molecular and regulatory mechanisms (Ruiz Castro et al., 2021). Previous transcriptomic studies in IBD have focused on gene-level differential expression associated with disease status (Song et al., 2025; Weiser et al., 2016; Kennedy Ng et al., 2026; Awad et al., 2026). Alternative splicing alterations have been reported in Crohn’s disease and ulcerative colitis individually (D. Li, Liang, et al., 2021a; D. Li and Tan, 2021), and in studies including both IBD subtypes (Häsler et al., 2016; Berger et al., 2021; Zhou et al., 2023). However, these studies have not directly addressed the extent to which splicing alterations contribute to transcriptomic differences between CD and UC.

In addition to characterizing genome-wide quantitative gene expression, RNA-seq data also capture the usage of specific exon–intron junctions, enabling systematic detection of alternative splicing events. Differential splicing (DS) is increasingly recognized as an important regulatory mechanism contributing to immune responses, cellular stress pathways, and disease pathogenesis (Jiang and Chen, 2021). Even in non-inflamed colonic mucosa, regulatory differences persist outside sites of active inflammation (Hong et al., 2017), providing a complementary setting in which to reveal baseline transcriptomic programs that distinguish CD and UC from non-IBD individuals beyond inflammation-driven changes and differences between these IBD subtypes. Integrating differential expression (DE) and DS offers a more complete view of transcriptomic regulation, as changes in gene-level abundance alone do not capture the extensive splicing variation that shapes cellular function.

Here, we performed a transcriptome-wide analysis integrating DE and DS analyses in macroscopically uninvolved colonic mucosal biopsies from 126 patients with CD, 99 patients with UC, and 71 non-IBD controls. We systematically characterized transcriptional alterations associated with each disease subtype compared to the non-IBD controls. DE and DS genes were largely non-overlapping in both CD and UC, indicating that these regulatory layers captured different components of IBD-associated mucosal dysregulation. CD and UC shared the majority of pathways enriched for either DS or DE genes including those related to chromatin regulation, RNA processing, and mRNA splicing. There were instances of pathway enrichments only seen in one of these comparisons, and directly comparing CD and UC patients revealed additional transcriptional differences related to both the splicing and expression of disease genes. In-depth analysis in two IBD-linked genes, *IRF1* and *CEACAM1*, demonstrated how altered splicing patterns can affect gene function that may contribute to disease. These results identified alternative splicing as a regulatory layer that informs on altered cellular processes distinct from gene-level expression analysis and that contributes to both shared and subtype-associated molecular programs in IBD.

## Results

### Differential splicing and expression distinguish IBD subtypes

We analyzed RNA-seq data from macroscopically uninvolved colonic mucosa of 126 CD patients, 99 UC patients, and 71 non-IBD subjects. In this study, we first separately compared CD and UC patients to the non-IBD cohort for both gene-level differential expression and intron-cluster-level differential splice site usage (Figure 1). Intron clusters contain two or more distinct splicing events involving multiple introns where a common set of splice junctions are shared. Genes may have multiple mutually-exclusive intron clusters.

In the CD vs. non-IBD comparison, we identified 520 differentially spliced (DS) intron clusters (cluster-adjusted *p <* 0.05, maximum absolute change in percent spliced in (*|*ΔPSI*|*) *≥* 0.029, the 90*^th^* percentile of *|*ΔPSI*|* in this comparison), corresponding to 491 unique DS genes (Supplementary Table 1). For UC patients, we found 1,451 significant DS clusters (cluster-adjusted *p <* 0.05, *|*ΔPSI*| ≥* 0.049), corresponding to 1,298 DS genes (Supplementary Table 2). Principal component analysis (PCA) of the residualized junction-count profiles showed good clustering of patient samples by their disease phenotypes (Figure 2a), indicating distinct splicing patterns characterized each group.

Next we examined disease-specific changes in gene expression. In CD patients, we identified 316 significant DE genes (*|*log_2_ fold change*| >* 1 and adjusted *p <* 0.05), including 269 genes upregulated and 47 genes downregulated compared to the non-IBD controls (Supplementary Table 3). In UC patients, we identified 1,107 significant DE genes, with 793 upregulated and 314 downregulated genes (Supplementary Table 4). PCA of residualized gene-expression profiles again showed separation by disease status, indicating gene expression could also distinguish sample groups (Figure 2b). Volcano plots (Figure 2c,d) reflected the greater number of DE genes that were upregulated than downregulated in disease in both comparisons. Overall, these results suggest that changes in splice site usage and expression levels are significant contributors to altered transcriptional programs in both CD and UC, but that there may also be differences across disease subtypes. In addition, more genes may be affected by alternative splicing than differential expression.

**Figure 1:**
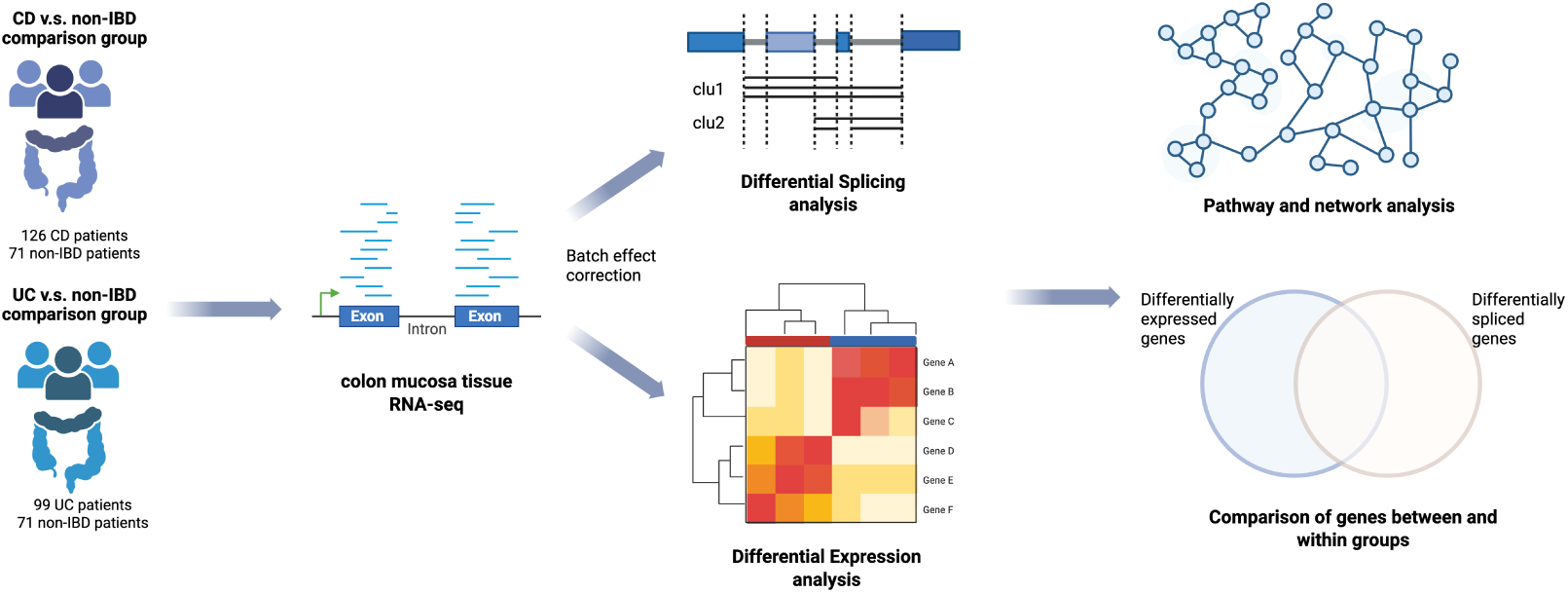
Study design. Colonic mucosal RNA-seq data from CD, UC, and non-IBD samples were processed and analyzed in parallel for differential expression and differential splicing. Significant DE and DS gene sets from CD vs. non-IBD and UC vs. non-IBD comparisons were compared within and across disease groups, along with pathway enrichment and network analyses.

**Figure 2:**
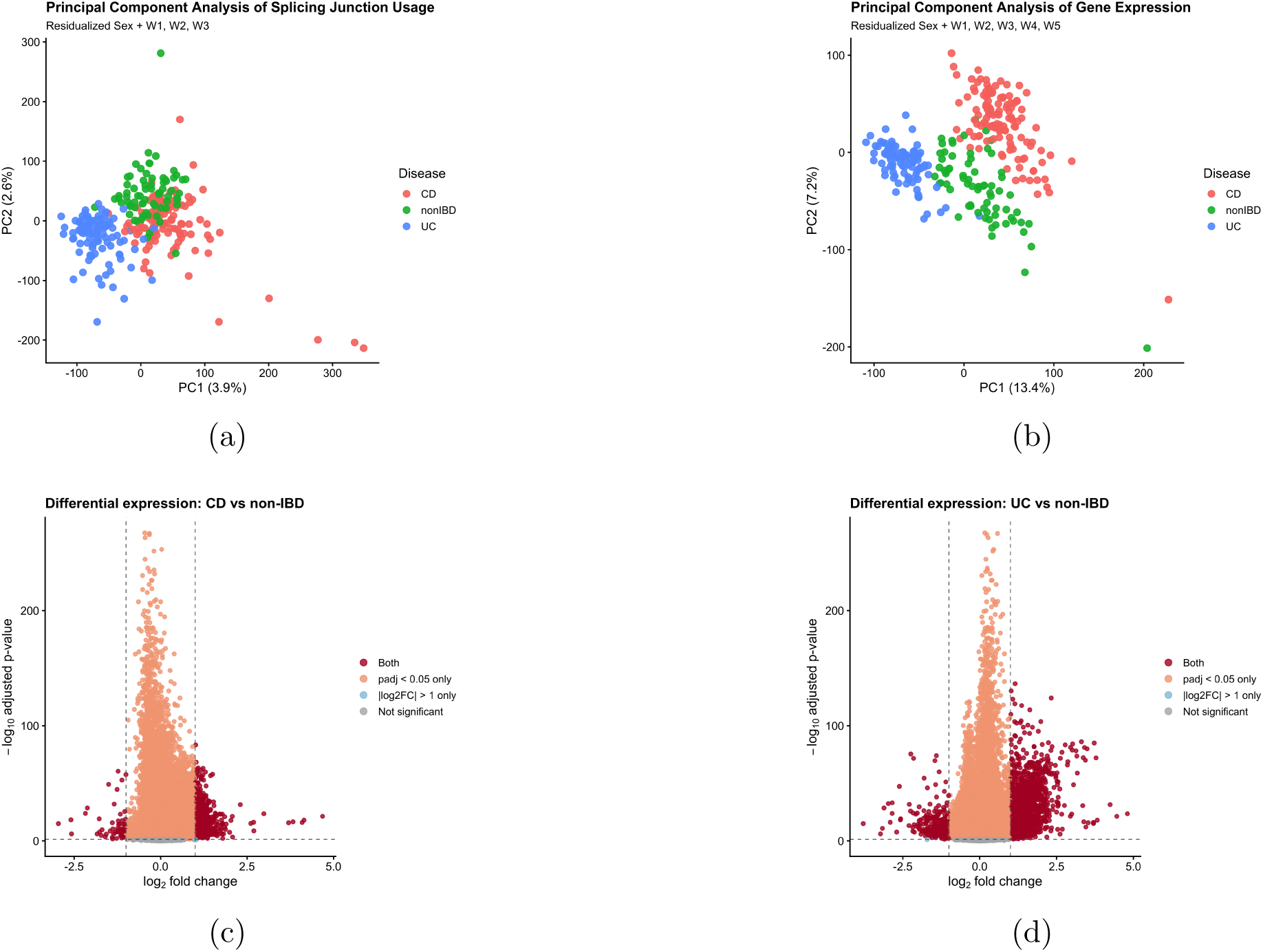
Global transcriptomic variation and disease-associated differential expression. **a)** Principal component analysis (PCA) of the residualized, normalized junction-count matrix, colored by disease status. Sex and the first three unwanted-variation factors (W1–W3), estimated with RUVSeq, were regressed out before visualization. PC1 and PC2 explained 3.9% and 2.6% of the total variance, respectively. **b)**PCA of the residualized, normalized gene-expression matrix, colored by disease status; sex and the first five RUVSeq unwanted-variation factors (W1–W5) were regressed out. PC1 and PC2 explained 13.4% and 7.2% of the total variance, respectively. In a–b, each point represents one sample. **c,d)** Volcano plots of differential expression for CD vs. non-IBD (**c**) and UC vs. non-IBD (**d**); each point represents one gene. Vertical dashed lines mark the effect-size threshold (*|*log_2_ fold change*| >* 1) and the horizontal dashed line the significance threshold (adjusted *p <* 0.05). Genes meeting both criteria (differentially expressed) are shown in dark red; genes meeting only the significance threshold in light red, only the effect-size threshold in light blue, and all others in grey.

### Differential splicing and expression identify largely distinct gene sets

We next determined whether a common set of genes showed both differential expression and splicing. In our analysis of CD patients, no genes were identified as DE and DS. In UC patients, only 16 genes were in common between the DE and DS genes, comprising 9 upregulated and 7 downregulated genes (Table 1). This limited overlap indicates that differential expression and differential splicing capture largely distinct disease-associated transcriptomic changes. Splicing analysis therefore reveals changes in transcript structure and intron usage that gene-level expression analysis alone would miss.

The 16 genes exhibiting both DE and DS in UC represent a small but biologically informative subset spanning several processes relevant to immune response, epithelial barrier function, and microbial interactions rather than converging on a single pathway. These include *IL1B*, a central pro-inflammatory cytokine implicated in UC-associated immune activation and anti-TNF nonresponse and central to immune–epithelial signaling (Aggeletopoulou et al., 2024; Zhang and Shi, 2020); *SGK1*, which is involved in epithelial ion transport and mucosal T-cell responses contributing to inflammatory signaling (Spagnuolo et al., 2018); *PLA2G2A*, a phospholipase induced in active colitis and proposed as a UC biomarker that is important for epithelial antimicrobial defense (Arijs et al., 2009; Ding et al., 2024); and *MUC12*, a transmembrane mucin that plays a key role in epithelial barrier and metabolic functions and was previously found to be altered in inflamed IBD mucosa (Niv, 2016; Leoncini et al., 2024; Grondin et al., 2020; Arras et al., 2025). Overall, these findings indicate that genes altered at both the expression and splicing levels in UC span diverse cellular processes and suggest coupled expression and splicing changes accompany multiple aspects of tissue function altered in UC.

**Table 1.** Genes exhibiting both differential expression and differential splicing in the UC vs. non-IBD comparison. log_2_ FC is the DESeq2 log_2_ fold change (UC relative to non-IBD) and DE_padj_ the corresponding Benjamini–Hochberg adjusted p-value. ΔPSI_max_ is the signed change in percent spliced-in of the intron with the largest absolute change within the gene’s most significant differentially spliced cluster (ΔPSI = PSI_UC_ *−* PSI_non-IBD_), and cluster_padj_ the adjusted p-value of that LeafCutter cluster. Differentially spliced clusters were defined as single-gene clusters with cluster-adjusted *p <* 0.05 and maximum *|*ΔPSI*| ≥* 0.049 (the 90th percentile of *|*ΔPSI*|* across all tested introns in the UC vs. non-IBD comparison). Gene descriptions are summarized from the National Center for Biotechnology Information (NCBI) Gene database.

| Gene | Description | $\log_2$<br>FC | $DE_{\text{padj}}$ | $\Delta\text{PSI}_{\text{max}}$ | $\text{Cluster}_{\text{padj}}$ |
| --- | --- | --- | --- | --- | --- |
| SLC30A10 | Manganese efflux transporter in epithelia. | 1.17 | $3.40 \times 10^{-14}$ | -0.085 | $3.70 \times 10^{-4}$ |
| APOBEC1 | Encodes a member of the cytidine deaminase enzyme family. | 1.24 | $1.40 \times 10^{-12}$ | -0.094 | 0.017 |
| CYP2B6 | Cytochrome P450 involved in drug metabolism and synthesis of cholesterol, steroids, and other lipids. | 1.04 | $1.23 \times 10^{-8}$ | 0.079 | 0.0233 |
| PITX2 | Transcription factor implicated in epithelial differentiation and Wnt pathways. | 1.03 | 0.0014 | -0.126 | $1.01 \times 10^{-5}$ |
| ANKRD36 | Predicted to enable ion channel inhibitor activity. | 1.09 | $1.54 \times 10^{-70}$ | 0.203 | $3.82 \times 10^{-3}$ |
| ABCC13 | Member of the gene superfamily encoding ATP-binding cassette (ABC) transporters. | 1.11 | $9.30 \times 10^{-43}$ | 0.096 | 0.00603 |
| HERC3 | Encodes a member of the HERC ubiquitin ligase family. | 1.42 | $9.34 \times 10^{-80}$ | -0.072 | 0.0094 |
| MGAM2 | Predicted to enable alpha-1,4-glucosidase activity. | 1.30 | $1.07 \times 10^{-26}$ | 0.069 | 0.0020 |
| LINC01612 | Long non-coding RNA. | 1.09 | $3.97 \times 10^{-14}$ | -0.127 | 0.0035 |
| PLA2G2A | Encodes a member of the phospholipase A2 (PLA2) family. | -1.34 | $2.97 \times 10^{-7}$ | 0.070 | $4.15 \times 10^{-13}$ |
| CKB | Encodes a cytoplasmic enzyme involved in cellular energy homeostasis. | -1.13 | $2.90 \times 10^{-12}$ | -0.060 | $3.65 \times 10^{-11}$ |
| MYH11 | Encodes a smooth muscle myosin heavy chain family protein. | -1.31 | $4.71 \times 10^{-10}$ | -0.096 | $1.46 \times 10^{-15}$ |
| IL1B | Encodes a member of the interleukin-1 cytokine family. | -1.05 | $1.01 \times 10^{-5}$ | 0.184 | 0.0023 |
| ACTG2 | Highly conserved protein involved in cell motility and cytoskeletal maintenance. | -1.83 | $1.11 \times 10^{-10}$ | 0.095 | $4.16 \times 10^{-17}$ |
| SGK1 | Encodes a serine/threonine protein kinase involved in stress response. | -1.06 | $2.83 \times 10^{-9}$ | 0.115 | $6.42 \times 10^{-6}$ |
| MUC12 | Encodes an integral membrane glycoprotein in the mucin family. | -1.46 | $1.62 \times 10^{-16}$ | 0.054 | $5.86 \times 10^{-9}$ |

### Differential expression and splicing reveal shared and disease-associated biological programs

To characterize the biological processes associated with altered expression and splicing in IBD, we performed pathway enrichment analysis for each of the CD vs. non-IBD and UC vs. non-IBD comparisons using Gene Ontology and Reactome annotations, with expressed genes as the enrichment background. For differential expression, separate analyses were performed for genes with increased vs. decreased expression in CD or UC. In total across all analyses, we identified 595 significantly enriched processes (adjusted *P <* 0.05), 579 from DE genes and 16 from DS genes. These corresponded to 383 distinct pathways, of which 239 were detected in at least one of the CD comparisons and 337 in at least one of the UC comparisons, with 193 shared (Supplementary Table 5). Because many terms represented overlapping gene sets, redundant terms were collapsed into 141 representative pathways (Supplementary Table 6). Pathway networks were constructed using the full sets of significantly enriched pathways from the three CD analyses (Figure 3) and three UC analyses (Figure 4). For more general interpretability, groups of related pathways were organized into modules and annotated according to the shared function of their member pathways.

As expected, both CD and UC DE analyses showed increased expression of genes in immune, signaling and antimicrobial defense pathways. Modules related to genome stability, DNA replication and chromatin regulation, including histone acetylation and heterochromatin organization, were also derived from genes with increased expression in both diseases (Figure 3, C1, C5, C6, C11 and C18; Figure 4, C1, C5, C7, C13 and C22). These common signals are consistent with previous evidence for altered transcriptional regulatory and epigenomic states across IBD. Profiling of human colonic biopsies has identified disease-associated changes in promoter and enhancer activity in both CD and UC, while integrated analyses of inflamed and non-inflamed mucosa have similarly demonstrated epithelial epigenomic alterations in both diseases (Ryan et al., 2020; Boyd et al., 2018). In the opposite direction, genes with decreased expression were enriched for pathways that formed a module involving keratan sulfate and glycosaminoglycan degradation in both diseases (Figure 3, C8; Figure 4, C10), which was the only downregulated module common to the two networks. Altered intestinal glycan biology is well established in IBD and can influence mucus integrity, host–microbial interactions and mucosal immune regulation (Kudelka et al., 2020). These shared modules indicate that CD and UC converge on several transcriptional programs in macroscopically uninvolved colonic mucosa despite their distinct clinical phenotypes.

In CD, while most modules were shared with the UC network, we detected four that were only significant in CD. Among genes with increased expression, these included antibody-mediated complement activation and Fc receptor signaling, consistent with previous reports of enhanced intestinal complement activity in CD (Preisker et al., 2019); triglyceride catabolism, in line with reported differences in lipid metabolism between CD and UC (Scoville et al., 2017); and broad infectious disease-related pathways. The remaining module, T-cell chemotaxis and migration, was derived from genes with decreased expression (Figure 3, Supplementary Table 7). Among DS genes, only two pathways were significantly enriched, extracellular exosome and extracellular vesicle (adjusted *P* = 0.049 for both), both of which were also enriched among genes with increased expression. However, these enrichments were driven by non-overlapping sets of DE and DS genes, indicating a combined contribution of expression and splicing to altered pathway function. The limited pathway enrichment of DS genes suggests that splicing alterations in CD were distributed across genes with diverse annotated functions rather than concentrated within specific pathways.

The UC network included a broader set of enriched epithelial and post-transcriptional programs than just those shared with CD (Figure 4, C6, C17, C21; Supplementary Table 8) and showed a more coherent pathway-level signature among DS genes than for CD. DS genes were significantly enriched for fourteen pathways, with the strongest enrichments involving mRNA binding, regulation of mRNA splicing, regulation of alternative splicing and RNA processing, which clustered into an mRNA-splicing and processing module (Figure 4, C29). The contributing DS genes included established RNA-processing factors including *PTBP1, RBFOX2, CELF1, TIA1, PRPF8 and SF3B1*. This indicates that differential splicing disproportionately affected genes involved in RNA processing itself, extending previous evidence of widespread alternative-splicing dysregulation in UC (D. Li and Tan, 2021) that occurs in the broader context of altered mucosal splicing reported across both major IBD subtypes (Häsler et al., 2016). A second module associated with DS genes involved COPII-mediated vesicle transport and ER-to-Golgi trafficking (Figure 4, C14), accompanied by significant enrichment of cytoplasmic stress-granule genes. The convergence of vesicular trafficking and stress-granule pathways is consistent with previous evidence of impaired epithelial stress responses and stress-granule formation in UC together with altered translation of proteins involved in secretion and epithelial homeostasis, even in histologically unaffected colonic mucosa (Tŕeton et al., 2011). Independent transcriptomic analysis of non-inflamed UC mucosa likewise identified altered ER-stress, glycosylation and vesicle-transport programs (Söderman, Berglind, and Almer, 2022). These observations provide independent support for the stress-response and secretory-trafficking components of the present enrichment, while the concentration of DS genes in RNA-processing pathways points to an additional post-transcriptional component of UC-associated mucosal dysregulation.

Across the UC disease network, enriched pathways for DE and DS genes were nearly distinct, with many more found for expression than splicing. No network edges connected DE- and DS-associated pathways, and no multi-pathway module contained both pathway types. In CD, only two terms were commonly enriched for both DE and DS genes but were driven by non-overlapping DE and DS gene sets. These reflect the general lack of overlap in DS and DE genes.

**Figure 3:**
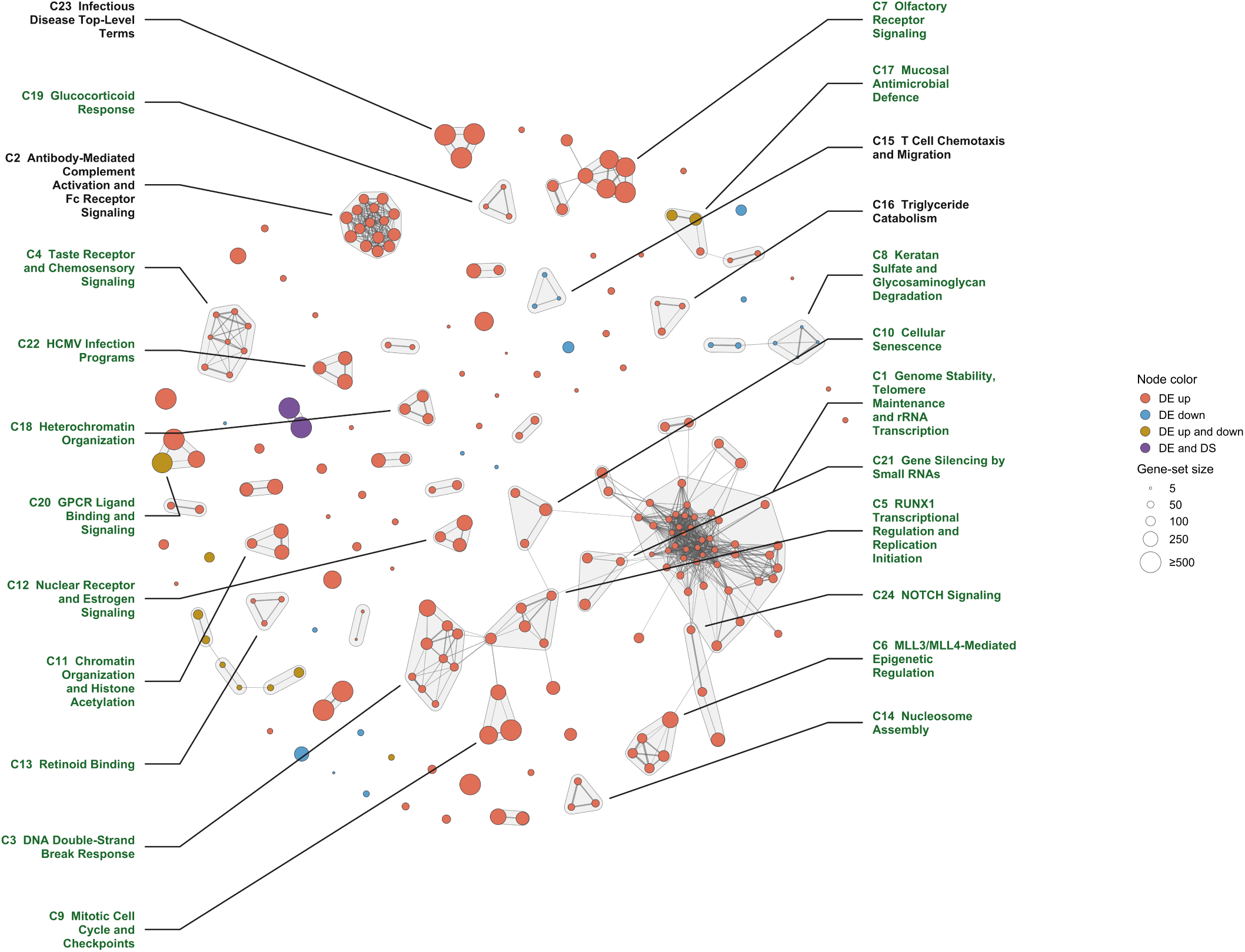
Functional network map of enriched pathways in the CD vs. non-IBD comparison. The network represents 239 pathways significantly enriched in the CD vs. non-IBD comparison (adjusted *p <* 0.05), using the expressed-gene background; GO 2026 and Reactome 2024). Enrichment was tested separately for upregulated DE, downregulated DE and DS genes. Nodes are colored according to the gene lists in which each pathway was enriched: orange, upregulated DE; blue, downregulated DE; yellow, both upregulated and downregulated DE; and purple, both DE and DS. Node size is proportional to pathway gene-set size and capped at 500 genes. Edges connect pathways with Jaccard similarity *≥* 0.35 between their gene sets, with edge width proportional to similarity. The network contains 742 such edges. Pathways were grouped into modules using Markov Cluster (MCL) clustering of the Jaccard-weighted network. Shaded outlines denote modules containing at least two pathways, whereas pathways not assigned to a multi-pathway module are shown as individual nodes. No module contained pathways enriched in both the upregulated and downregulated DE gene lists. Modules were manually annotated according to shared biological functions, and only the 24 modules containing at least three pathways are labeled. The leader lines connect labels to the corresponding modules. Green labels denote modules also identified in the UC network, defined by *≥* 50% member-pathway overlap, whereas black labels denote the four modules specific to the CD network. Pathways comprising each labeled module are listed in Supplementary Table 7.

**Figure 4:**
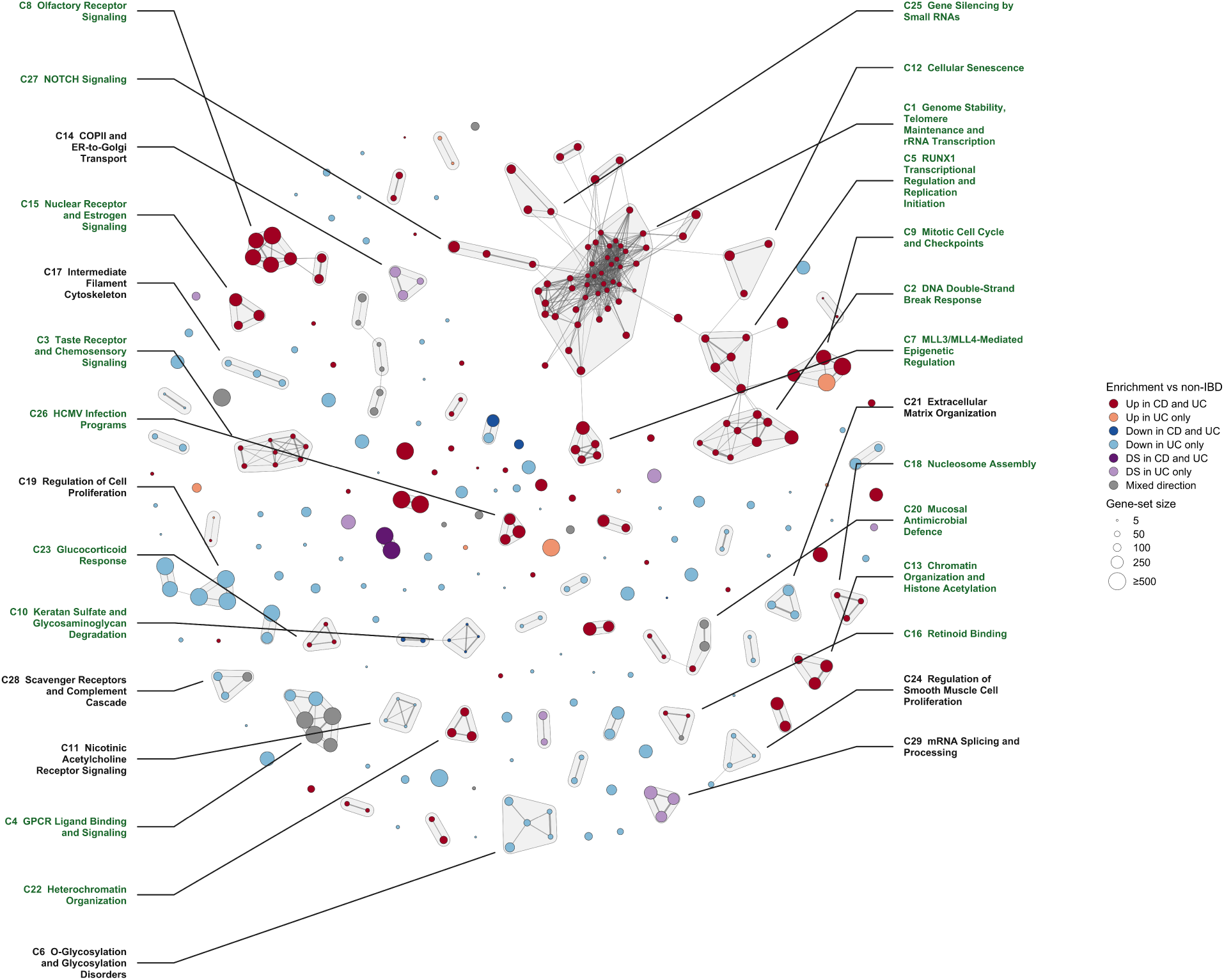
Functional network map of enriched pathways in the UC vs. non-IBD comparison. The network represents 337 pathways significantly enriched in the UC vs. non-IBD comparison (adjusted *p <* 0.05, using the expressed-gene background; GO 2026 and Reactome 2024). Enrichment was tested separately for upregulated DE, downregulated DE and DS genes. Nodes are colored according to the direction of enrichment and whether the corresponding pathway was also enriched in CD vs. non-IBD: dark red, upregulated DE in both CD and UC; orange, upregulated DE in UC only; dark blue, downregulated DE in both CD and UC; light blue, downregulated DE in UC only; dark purple, DS in both CD and UC; light purple, DS in UC only; and grey, mixed enrichment directions. Within each regulatory category, darker colors distinguish pathways enriched in both diseases from those enriched only in UC. Node size is proportional to pathway gene-set size and capped at 500 genes. Edges connect pathways with Jaccard similarity *≥* 0.35 between their gene sets, with edge width proportional to similarity. The network contains 685 such edges. Pathways were grouped into modules using Markov Cluster (MCL) clustering of the Jaccard-weighted network. Shaded outlines denote modules containing at least two pathways, whereas pathways not assigned to a multi-pathway module are shown as individual nodes. No module contained pathways enriched among both upregulated and downregulated DE genes, and no module contained both DE- and DS-associated pathways. Modules were manually annotated according to the shared biological functions of their member pathways, and only the 29 modules containing at least three pathways are labeled. Leader lines connect labels to their corresponding modules. Green labels denote modules also identified in the CD network, defined by *≥* 50% member-pathway overlap, whereas black labels denote the nine modules identified only in the UC network. Pathways comprising each labeled module are listed in Supplementary Table 8.

### Additional disease subtype-preferential altered programs identified by direct comparison of CD and UC patients

Pathways identified in separate disease vs. control comparisons do not necessarily allow us to identify significant transcriptional differences between CD and UC patients. We therefore compared the two disease subtypes directly using the same analytical pipelines, significance thresholds and expressed-gene background. This CD vs. UC comparison identified 187 genes expressed significantly higher in UC, 313 expressed significantly higher in CD (Supplementary Table 9) and 1,520 differentially spliced genes, of which 269 DE and 930 DS genes were also significant in the UC vs. non-IBD comparison and 43 DE and 206 DS genes in the CD vs. non-IBD comparison (Supplementary Table 10).

Pathway enrichment analysis of DE and DS genes in the direct CD vs. UC comparison identified 147 significant enrichment results (adjusted *P <* 0.05), comprising 145 distinct pathways. Of these, 16 pathways were associated with genes expressed more highly in UC, 116 with genes expressed more highly in CD and 15 with DS genes. Comparison with the disease vs. control analyses showed that among the 145 distinct pathways, 77 pathways were identified only in this direct comparison, 36 had previously been enriched only in UC vs. non-IBD, 16 only in CD vs. non-IBD and 16 in both disease vs. control comparisons (Figure 5, Supplementary Table 11).

At the expression level, most pathways also seen in both disease vs. control comparisons are enriched among genes expressed more highly in UC and were related to taste receptor, chemosensory and G protein-coupled receptor signaling. Of these, 12 had also been enriched among genes upregulated in both CD and UC relative to non-IBD controls, forming chemosensory signaling modules in both networks (Figure 3, C4, and Figure 4, C3). Their enrichment in the direct comparison suggests a more pronounced chemosensory expression signature in UC despite increased expression in both diseases. Taste receptors are expressed throughout the gastrointestinal mucosa, including enteroendocrine, goblet, Paneth and tuft cells, where they participate in luminal sensing, antimicrobial defense and mucosal immune responses (Sternini and Rozengurt, 2024). Recent analysis of colonic biopsies from patients with active UC also reported altered expression of several mucosal taste receptors relative to healthy controls, providing independent evidence that intestinal chemosensory signalling is perturbed in UC (Y. Liu et al., 2025).

In contrast, genes expressed more highly in CD were predominantly associated with humoral immune responses, including complement activation, Fc receptor signaling and B-cell receptor pathways, consistent with pathways previously enriched in the CD vs. non-IBD comparison. Of the 313 genes expressed more highly in CD, 65 were immunoglobulin genes, including 58 variable-region segments, which largely accounted for 39 of the 116 enriched pathways. In addition to this immunoglobulin signature, genes expressed more highly in CD included many complement-related genes. This result is consistent with previous direct comparisons of intestinal complement activity in CD and UC, reporting higher mucosal C1q and C3 abundance in CD than UC and increased expression of multiple complement components in CD, including in non-inflamed mucosa during remission (Preisker et al., 2019).

For genes with differential splicing usage, we identified three RNA processing pathways that were also enriched in UC DS genes when compared to controls, indicating that part of the UC post-transcriptional signature distinguished the two subtypes. Interestingly, the remaining pathways were nearly all unique to this direct disease subtype comparison and seemed related to epithelial barrier function. One module involved pathways related to epithelial cell structure and adhesion. Previous studies showed that disruption of epithelial RhoA signalling or *β*-actin compromises cytoskeletal organization, cell shedding and barrier integrity, and increases susceptibility to intestinal inflammation (López-Posadas et al., 2016; Lechuga et al., 2020). A second module involved genes related to Golgi-associated protein processing, including glycosylation. Glycosylation related genes included *MPI* and other genes involved in glycoprotein maturation and secretory trafficking. Experimental loss of epithelial *MPI* was shown to impair N-glycosylation, disrupt MUC2 maturation and secretion, induce ER stress and compromise the mucus barrier (Roy et al., 2026). Since DS enrichment is not directional, it is unclear how these altered splicing programs may differentially affect the two diseases. Overall, these analyses point to processes that may be perturbed in CD and UC in different ways, contributing to the distinct phenotypes seen in these IBD subtypes.

**Figure 5:**
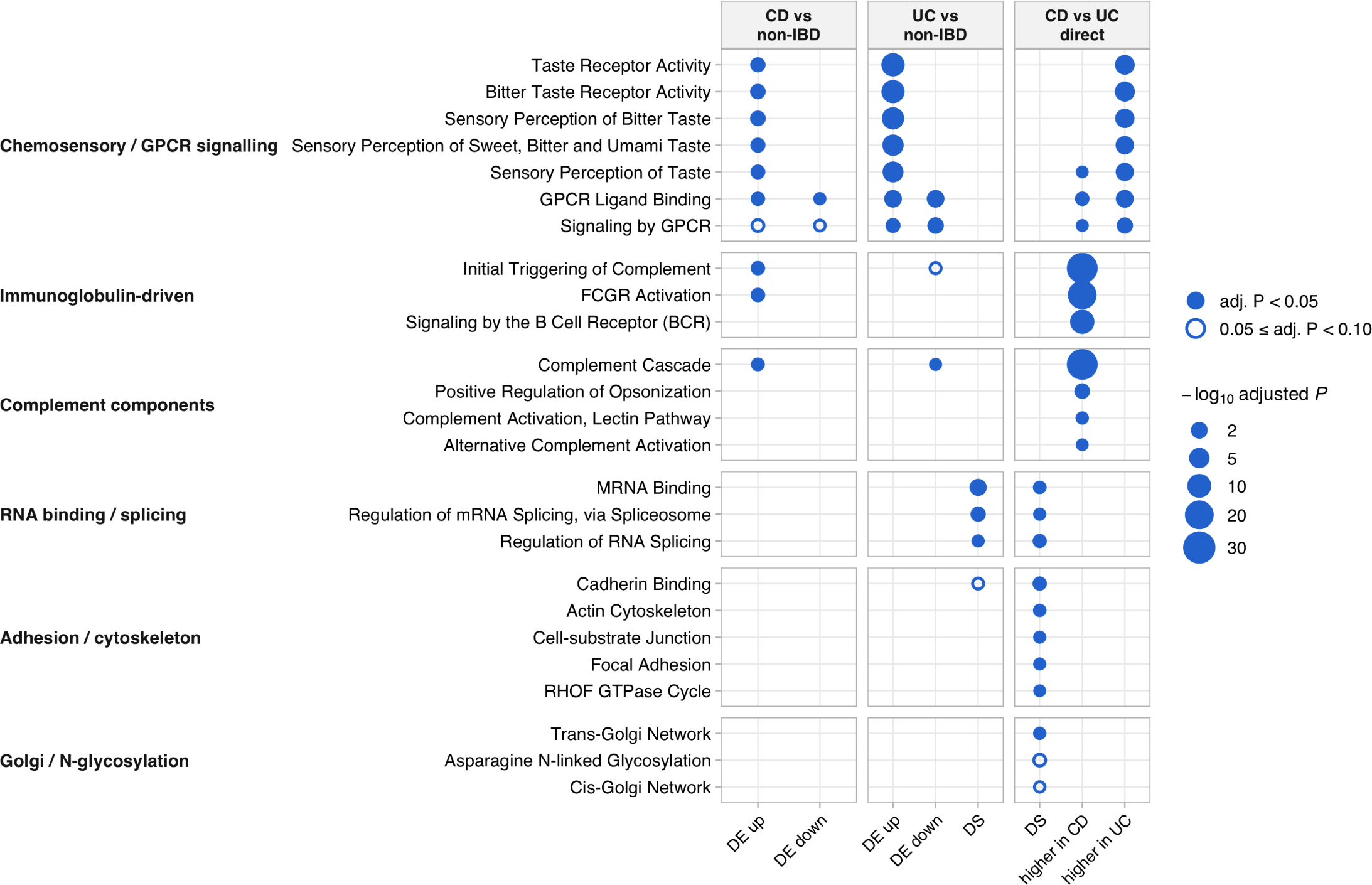
Pathway enrichment across disease vs. control and direct subtype comparisons. Pathways discussed in the text are grouped by biological theme (rows) and shown across the CD vs. non-IBD, UC vs. non-IBD and direct CD vs. UC comparisons (columns). Within each comparison, pathways were tested using DE and DS gene sets against the expressed-gene background. For disease vs. control comparisons, DE genes were separated into upregulated and downregulated sets; for the direct comparison, they were separated into genes expressed more highly in CD or UC. Filled circles indicate pathways with an adjusted *p <* 0.05 and open circles pathways with an adjusted *p <* 0.10, with circle size proportional to *—* log_10_(adjusted *p*). Columns without a significant pathway are omitted. One Reactome pathway name is abbreviated. Complete enrichment results are provided in Supplementary Tables 5 and 11.

### Differential splicing at *IRF1* and *CEACAM1* may shift aspects of their function that contribute to disease

Having characterized differential expression and splicing transcriptome-wide, we examined two genes in detail to demonstrate how changes in splicing usage may affect key cellular processes in which these genes play an important role. *IRF1* and *CEACAM1* were each differentially spliced without meeting the threshold for differential expression (Table 2), indicating that changes in junction usage did not coincide with detectable changes in total gene abundance. IRF1 is an immune-regulatory transcription factor involved in inflammatory signaling, whereas CEACAM1 regulates epithelial and mucosal immune functions (Tang et al., 2014; Berger et al., 2021). We therefore investigated whether the observed splicing differences could affect transcript regions relevant to their respective functions.

**Table 2.** Representative genes exhibiting differential splicing without differential expression. Each gene represents a DS-only example that met the differential-splicing criteria in the indicated comparison but did not meet the differential-expression criteria. Differential splicing was defined by a cluster-level adjusted *p <* 0.05 and a maximum *|*ΔPSI*|* meeting or exceeding the 90th percentile of *|*ΔPSI*|* across all tested introns in the corresponding comparison (0.029 for CD vs. non-IBD and 0.049 for UC vs. non-IBD). Differential expression was defined as *|*log_2_ FC*| >* 1 and adjusted *p <* 0.05. For each gene, all introns belonging to the representative significant cluster are shown and ordered by the magnitude of ΔPSI. PSI_non-IBD_ and PSI_disease_ denote the fraction of cluster-level spliced reads supporting each intron in the respective groups. ΔPSI was calculated as PSI_disease_ *−* PSI_non-IBD_, such that positive values indicate increased intron usage in disease. cluster_padj_ denotes the Benjamini–Hochberg-adjusted *p* value from the LeafCutter cluster-level likelihood-ratio test and is reported once per cluster.

| Gene | Comparison | Intron | PSI<sub>non-IBD</sub> | PSI<sub>disease</sub> | ΔPSI | cluster<sub>padj</sub> |
| --- | --- | --- | --- | --- | --- | --- |
| IRF1 | CD vs. non-IBD | chr5:<br>132487130–132487926 | 0.4283 | 0.4725 | 0.0441 | $9.86 \times 10^{-4}$ |
|  |  | chr5:<br>132489483–132490545 | 0.2776 | 0.2545 | −0.0231 |  |
|  |  | chr5:<br>132488025–132489392 | 0.2894 | 0.2681 | −0.0213 |  |
|  |  | chr5:<br>132488025–132490545 | 0.0022 | 0.0024 | 0.0002 |  |
|  |  | chr5:<br>132487130–132490545 | 0.0025 | 0.0025 | 0.0001 |  |
| CEACAM1 | UC vs. non-IBD | chr19_clu9999:<br>42519235–42521267 | 0.5093 | 0.4176 | −0.0917 | $4.68 \times 10^{-50}$ |
|  |  | chr19_clu9999:<br>42512479–42518948 | 0.4477 | 0.5387 | 0.0910 |  |
|  |  | chr19_clu9999:<br>42512479–42521267 | 0.0294 | 0.0325 | 0.0031 |  |
|  |  | chr19_clu9999:<br>42512479–42516773 | 0.0086 | 0.0070 | −0.0016 |  |
|  |  | chr19_clu9999:<br>42519101–42521267 | 0.0050 | 0.0041 | −0.0009 |  |
| | | chr19_clu9998:<br>42510920–42512350 | 0.7009 | 0.7428 | 0.0419 | $1.17 \times 10^{-23}$ |
|  |  | chr19_clu9998:<br>42510920–42511576 | 0.1574 | 0.1198 | −0.0376 |  |
|  |  | chr19_clu9998<br>42511628–42512350 | 0.1147 | 0.0898 | −0.0249 |  |
|  |  | chr19_clu9998:<br>42510915–42512350 | 0.0270 | 0.0476 | 0.0206 |  |

| Gene | Comparison | Intron | PSI<sub>non-IBD</sub> | PSI<sub>disease</sub> | ΔPSI | cluster<sub>padj</sub> |
| --- | --- | --- | --- | --- | --- | --- |

*IRF1* is an inflammation-induced transcription factor that regulates immune responses and intestinal epithelial cell survival and has been implicated in TNF-*α*-driven epithelial injury in IBD (Tang et al., 2014). In our analysis, *IRF1* exhibited differential splicing in CD without differential expression. The corresponding intron cluster was not tested in the UC vs. non-IBD comparison because dataset-specific LeafCutter clustering grouped these junctions with distant, low-coverage junctions, leading to their removal during filtering. Similarly, in the direct CD vs. UC comparison, LeafCutter constructed a different cluster retaining only three of the original five junctions. This cluster was not significant (adjusted *p* = 0.37), and the retained junctions showed similar relative usage between CD and UC. Thus, although *IRF1* was differentially spliced in CD relative to non-IBD controls, these results do not confidently indicate that the event is specific to CD. A previous study reported altered *IRF1* transcript usage in colonic tissue, including an increased proportion of transcripts with an extended 3*^′^* untranslated region (D. Li, Liang, et al., 2021b). In contrast, the cluster identified here spanned exons 1–4 at the 5*^′^* end of the gene (Figure 6a). Exon 1 (E1) is completely contained in the 5’ untranslated region, while exon 2 (E2) contains the translation initiation codon and part of the DNA-binding domain. Transcripts lacking E2 can be translated only into N-terminally truncated proteins that lack DNA-binding activity and fail to induce expression of IRF1 target genes (Maratheftis et al., 2006). In our data, the canonical exon 3 to exon 4 (E3–E4) junction usage increased in CD (ΔPSI = +0.044), whereas usage of the two junctions flanking E2, E1–E2 and E2–E3, decreased (ΔPSI = *−*0.023 and *−*0.021, respectively; Figure 6a,b). However, junctions that skip E2 by joining E1 directly to E3 or E4 (E1–E3 and E1–E4) remained rare in both groups (PSI *≤* 0.003), indicating that the shift does not reflect increased E2 skipping. We therefore hypothesised that it reflects changes in expression of transcripts that begin downstream of E2 and contain E3–E4 but neither upstream junction. Of the 21 annotated *IRF1* transcripts containing at least one junction from this cluster, two have this structure: ENST00000472045, a retained-intron transcript that begins within intron 2, and ENST00000681462, a transcript annotated for nonsense-mediated decay that begins at exon 3 (Figure 6a). Both lack E2 and would therefore not produce a DNA-binding form of IRF1. Because junction-level data cannot establish which transcript structures give rise to this pattern, we performed long-read sequencing on a small number of samples to assess isoform transcript expression in colonic mucosa to determine which transcript isoforms were present.

The long-read data comprised two non-IBD, two CD and two inflamed UC biopsies. Nine annotated *IRF1* transcripts had assigned reads in at least one sample (Figure 6c). Most reads were assigned to the two full-length transcripts ENST00000245414 and ENST00001132324, and transcripts containing all three canonical junctions of the cluster accounted for 81–94% of *IRF1* -assigned reads per sample. ENST00000472045, for which transcription begins within the canonical intron 2 and lacks the DNA binding domain, was detected in four of six samples (Figure 6c). Most of the reads assigned to this transcript were compatible only with this transcript, indicating that its detection did not depend on probabilistic assignment of ambiguous reads. ENST00000681462 had no assigned reads and of the two annotated E2-skipping transcripts, ENST00000679921 was detected at low abundance, and ENST00000680352 was not detected (Figure 6c). These data confirm that an *IRF1* transcript beginning downstream of E2 is expressed in colonic mucosa and that annotated E2-skipping transcripts are uncommon, consistent with the rarity of the E1–E3 and E1–E4 junctions. While differential abundance of splicing events in CD could not be confirmed with these few samples, they support the possibility that the detected differential splicing usage could indicate reduced transcription of canonical transcripts and increased transcription of an isoform lacking the DNA binding domain leading to reduced induction of *IRF1* target genes in CD.

**Figure 6:**
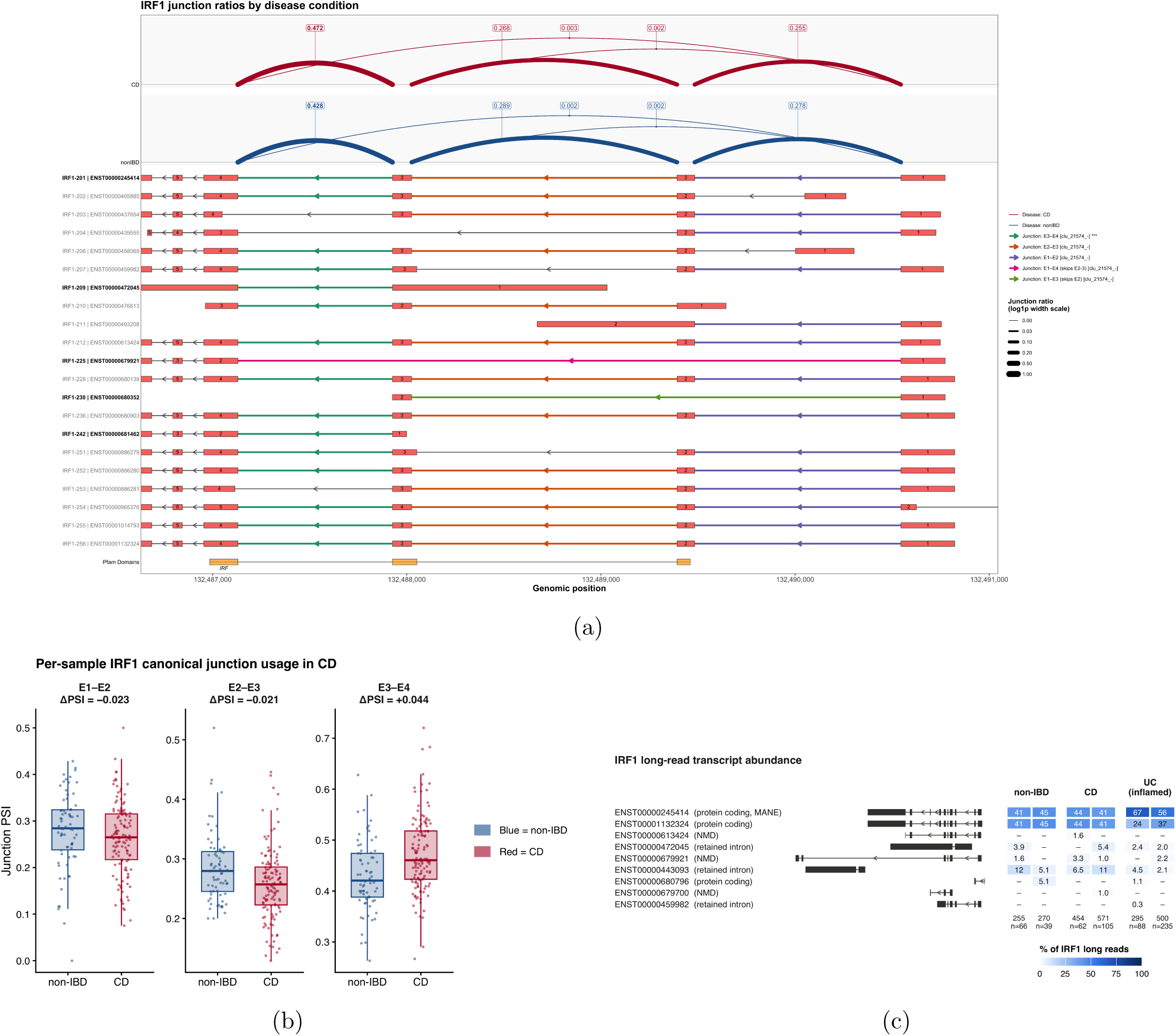
Differential junction usage in *IRF1* in CD. **(a)** LeafCutter junction ratios for chr5_clu21574 (cluster-adjusted *P* = 9.9 *×* 10^−4^; Table 2), shown above the 21 GENCODE v50 *IRF1* transcripts containing at least one cluster junction. CD is shown in red and non-IBD in blue. Arc labels indicate fitted mean junction PSI; arc width is scaled logarithmically with PSI. Cluster junctions are highlighted in the transcript models, and the bottom track indicates the Pfam DNA-binding domain, which begins in exon 2. Transcript labels highlighted in bold black indicate isoforms discussed in the text, whereas other annotated transcripts are labeled in grey. Exon numbers refer to positions within each transcript. Genomic coordinates increase from left to right. *IRF1* is on the minus strand, and transcription proceeds from right to left. **(b)** Per-sample PSI for the three junctions in samples with at least 20 cluster-level reads. Boxes indicate medians and interquartile ranges while whiskers extend to 1.5*×* the interquartile range, and points denote individual samples. ΔPSI = PSI_CD_ *−* PSI_non-IBD_ is estimated by LeafCutter. **(c)** Oxford Nanopore reads assigned by bambu to annotated *IRF1* transcripts in two non-IBD, two CD, and two inflamed UC biopsies. Left, GENCODE v50 structures of the transcripts with assigned reads. Exon widths are drawn to scale, whereas introns are compressed to a uniform width. *IRF1* is located on the minus strand, with transcription proceeding from right to left (arrowheads). Right, percentage of all *IRF1* -assigned reads attributed to each transcript in each sample. *n*, total number of *IRF1* -assigned reads per sample; –, no assigned reads. PSI, proportion of reads supporting a given junction among all reads assigned to its LeafCutter intron cluster.

*CEACAM1* is an established IBD risk gene that regulates mucosal immune responses, including those mediated by intestinal T cells (Berger et al., 2021). Extensive alternative splicing generates isoforms that differ in their extracellular and cytoplasmic domains, primarily through alternative inclusion of exon 5 (E5), which encodes the extracellular A2 immunoglobulin-like domain, and exon 7 (E7), which determines cytoplasmic tail length (Helfrich and Singer, 2019; Dery et al., 2024). E5 skipping produces isoforms lacking the A2 domain, whereas E7 inclusion generates the long-tail isoform (CEACAM1-L), which contains two immunoreceptor tyrosine-based inhibitory motifs. Conversely, E7 skipping produces the short-tail isoform (CEACAM1-S), which lacks these motifs (Helfrich and Singer, 2019; Dery et al., 2024).

In our analysis, *CEACAM1* exhibited differential splicing in UC without differential expression. The same intron cluster was also significant at the cluster level in CD vs. non-IBD (adjusted *p* = 0.012) but did not meet our effect-size criterion (maximum *|*ΔPSI*|* = 0.022). In the direct CD vs. UC comparison, the same five-junction cluster was significantly differentially spliced (adjusted *p* = 6.2 *×* 10^−49^, maximum *|*ΔPSI*|* = 0.071), providing evidence of a subtype-associated difference in junction usage. Previous studies have identified isoforms lacking the E5-encoded A2 domain in advanced melanoma and showed that their expression can be induced by interferons and the TLR3 agonist poly(I:C) in human bronchial cells (Dery et al., 2024). In the intron cluster spanning exons 4–6 (E4–E6) with significantly altered usage patterns in UC compared to controls, E4–E5 junction usage decreased whereas E5–E6 usage increased (ΔPSI = *−*0.092 and +0.091, respectively; Table 2; Figure 7a,b). This same pattern was observed in the comparison with CD (E4–E5 reduced: ΔPSI = *−*0.069; E5–E6 increased: ΔPSI = +0.071). However, the E4–E6 junction, which skips E5, showed little change (ΔPSI = +0.003), indicating that the redistribution of intron usage was not driven by increased canonical E5 skipping. Instead, the opposing changes in usage of the two junctions could reflect altered transcription of isoforms containing only one of these junctions. Candidate annotated transcripts include ENST00000485605, where transcription initiates within the canonical intron 4 and contains E5–E6 but not E4–E5; ENST00000979359 and ENST00000979362, which use an alternative splice acceptor within E5; and ENST00000377806 and ENST00000867318, which contain E4–E5 but not E5– E6 (Figure 7a). Several are annotated as retained-intron or nonsense-mediated-decay transcripts. To investigate whether these alternative structures are present in colonic mucosa, we again examined our complementary long-read sequencing data.

Nineteen annotated *CEACAM1* transcripts had evidence of expression in the long-read data in at least one sample (Figure 7c). Transcripts containing both E4–E5 and E5–E6 accounted for most assigned reads (59–78% per sample), whereas transcripts skipping E5 represented only 2.5–3.6%. Transcripts containing E5–E6 without E4–E5 were also detected, although 97% of the reads in this class were assigned to a single transcript, ENST00000979359, of which only two reads were uniquely compatible with that structure. Its estimated abundance therefore depended largely on probabilistic assignment of reads compatible with multiple isoforms. By contrast, nine of the ten reads assigned to ENST00000485605 were uniquely compatible with its annotated structure, providing more specific evidence of its expression, although it represented only 0.4% of assigned reads. While these data support the expression of transcripts containing only one canonical junction, they cannot establish exactly which isoforms account for the UC-associated redistribution.

A second intron cluster spanning exons 6–8 (E6–E8) showed increased E6–E8 junction usage, accompanied by decreased E6–E7 and E7–E8 usage, in UC (Figure 7a,b). Although statistically significant (adjusted *p* = 1.2 *×* 10^−23^), the cluster did not meet our differential-splicing criteria because its maximum *|*ΔPSI*|* of 0.042 was below the prespecified effect-size cutoff of 0.049 for UC vs. non-IBD. Similarly, in the UC vs. CD comparison, this was statistically significant (adjusted *p* = 3 *×* 10^−25^) but with a *|*ΔPSI*|* of 0.046 that again missed the effect-size cutoff. We nevertheless examined this cluster because E7 inclusion determines the cytoplasmic tail of CEACAM1. Previous studies have demonstrated that E7 splicing is regulated by specific RNA-binding proteins and that IRF1 promotes CEACAM1-L production in breast epithelial cell models (Dery et al., 2024). The balance between these isoforms has also been associated with intestinal disease phenotypes. In colorectal cancer, CEACAM1-L predominance at the invasive front was associated with lymph-node and haematogenous metastasis and shorter survival, while cell-based experiments showed that CEACAM1-L promoted invasion and migration (Ieda et al., 2011). In IBD, Berger et al. reported reduced E7 inclusion in ileal biopsies from two patients with CD exhibiting aberrant, rectal-like splicing profiles (Berger et al., 2021). The direction of this previously reported change is consistent with the increased E6–E8 junction usage observed here in UC, although the findings concern different disease subtypes and intestinal regions.

Long-read sequencing supported the expression of both E7-containing and E7-skipping transcript structures in colonic mucosa, including ENST00000161559 and ENST00000352591 that include E7, and ENST00000979360 and ENST00000485605 that exclude it (Figure 7c). However, most reads were compatible with multiple annotated transcripts, limiting the reliability of isoform-specific abundance estimates. Thus, the long read data provides evidence for the expression both types of transcripts, making it possible that there is increased E7 exclusion in UC as suggested in the short-read junction estimates.

**Figure 7:**
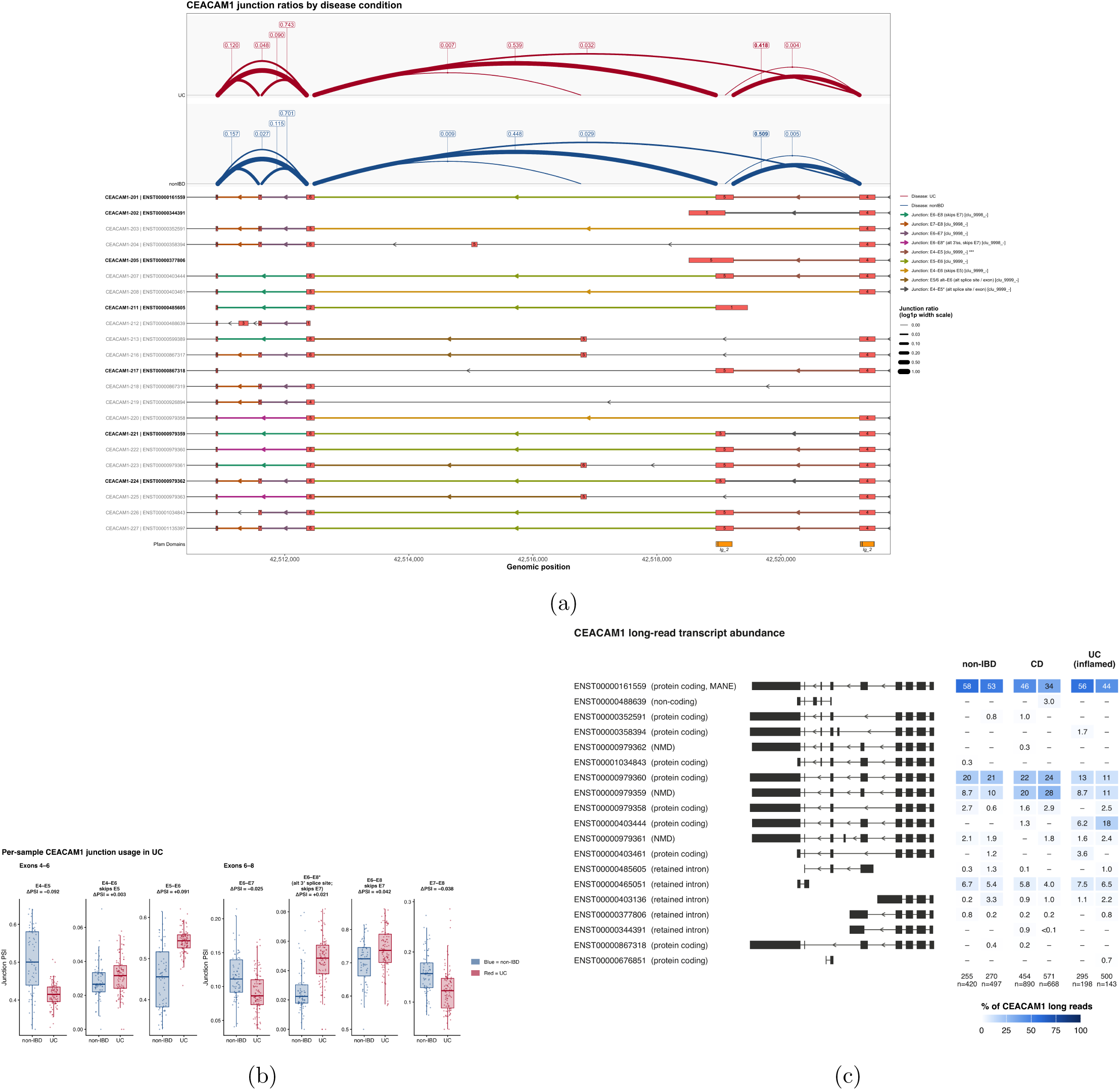
Junction usage and transcript structure in *CEACAM1* in UC. **(a)** LeafCutter junction ratios for chr19_clu9999 (exons 4–6; cluster-adjusted *p* = 4.7 *×* 10^−50^; Table 2) and chr19_clu9998 (exons 6–8; cluster-adjusted *p* = 1.2 *×* 10^−23^, maximum *|*ΔPSI*|* = 0.042), shown above 22 GENCODE v50 *CEACAM1* transcripts containing at least one cluster junction. UC is shown in red and non-IBD in blue. Arc labels indicate fitted mean junction PSI; arc width is scaled logarithmically with PSI. Cluster junctions are highlighted in the transcript models. Asterisks mark alternative splice acceptors (E4–E5*^∗^*, 134 bp within exon 5; E6–E8*^∗^*, 5 bp within exon 8). The bottom track indicates the Pfam Ig_2 domains encoded by exons 4 and 5. Transcript labels highlighted in bold black indicate isoforms discussed in the text, whereas other annotated transcripts are labeled in grey. Exon numbers refer to positions within each transcript. Genomic coordinates increase from left to right. *CEACAM1* is on the minus strand, transcription proceeds from right to left. **(b)** Per-sample PSI for junctions in both clusters in samples with at least 20 cluster-level reads. Boxes indicate medians and interquartile ranges while whiskers extend to 1.5*×* the interquartile range, and points denote individual samples. ΔPSI = PSI_UC_ *−* PSI_non-IBD_ is estimated by LeafCutter. **(c)** Oxford Nanopore reads assigned by bambu to annotated *CEACAM1* transcripts in two non-IBD, two CD, and two inflamed UC biopsies. Left, GENCODE v50 structures of the transcripts with assigned reads. Exon widths are drawn to scale, whereas introns are compressed to a uniform width. *CEACAM1* is located on the minus strand, with transcription proceeding from right to left (arrowheads). Right, percentage of all *CEACAM1* -assigned reads attributed to each transcript in each sample. *n*, total number1o9f *CEACAM1* -assigned reads per sample; –, no assigned reads. PSI, proportion of cluster-level spliced reads supporting a given junction.

## Discussion

In this study, we integrated differential expression and differential splicing analyses in macroscopically uninvolved colonic mucosa from 126 CD patients, 99 UC patients and 71 non-IBD controls. We identified 316 DE and 491 DS genes in CD, and 1,107 DE and 1,298 DS genes in UC compared with non-IBD controls. Despite these substantial numbers, expression and splicing changes showed little overlap: no genes were both DE and DS in CD, and only 16 overlapped in UC. By examining these regulatory layers in parallel and directly comparing CD with UC, our study identifies disease-associated splicing alterations and additional molecular differences between the two IBD subtypes that would not be detected through gene-level expression analysis alone.

In contrast to the broad functional enrichment observed among DE genes, the extensive differential splicing identified in both diseases was accompanied by relatively restricted pathway enrichment. This pattern is consistent with previous observations that expression and splicing changes often affect genes with distinct biological functions (Dam, Olsen, and Vitting-Seerup, 2023; Häsler et al., 2016), but raises questions about how the biological consequences of widespread splicing alterations should be interpreted. One possibility is that splicing changes are distributed across genes with diverse functions rather than concentrated within coordinated biological programs, as also observed across other human transcriptomic datasets (Dam, Olsen, and Vitting-Seerup, 2023). Alternatively, conventional pathway analyses may not adequately capture isoform-specific changes, as gene-level functional annotations do not distinguish transcripts with different protein domains or regulatory properties. Thus, limited pathway enrichment does not necessarily imply limited biological relevance. The enrichment of RNA-processing and splicing-regulatory genes among UC DS genes is particularly interesting in this context. Because these factors regulate the processing of other transcripts, changes in their own splicing could potentially influence broader post-transcriptional regulation. The accompanying enrichment of stress-granule and secretory-trafficking pathways also places these alterations in the context of persistent stress-response and secretory abnormalities previously reported in non-inflamed UC mucosa (Tréton et al., 2011; Söderman, Berglind, and Almer, 2022). This suggests that altered transcript processing contributes to broader regulatory changes in UC, although the functional relationships between these processes remain unclear.

Our direct comparison of CD and UC revealed subtype-associated differences that were not apparent from the separate disease vs. control analyses. Although several expression-associated pathways were enriched in both diseases relative to controls, only a subset significantly distinguished CD from UC, indicating that disease-associated changes do not necessarily represent differences between the subtypes. The stronger chemosensory expression signature in UC and humoral immune and complement-associated signatures in CD suggest differences in mucosal environmental sensing and immune regulation. Notably, the direct splicing comparison identified shifts in splice-site usage in genes associated with epithelial adhesion, cytoskeletal organization and Golgi-associated protein processing, several of which were not enriched in the separate disease vs. control comparisons. These processes are important for epithelial barrier maintenance, and experimental studies have demonstrated that disruption of cytoskeletal regulation or glycoprotein maturation can compromise intestinal epithelial integrity and increase susceptibility to inflammation (Lechuga et al., 2020; Roy et al., 2026). Our findings therefore indicate that CD and UC differ not only in immune-associated gene expression but also in transcript processing within pathways supporting epithelial structure and secretory function. Specifically how these shifts might contribute to phenotype differences in the IBD subtypes warrants further investigation.

The *IRF1* and *CEACAM1* analyses illustrate the potential biological significance of the DS-only genes in CD and UC. Both exhibited altered junction usage involving transcript regions relevant to protein function, and previous studies have shown that truncated IRF1 isoforms can lose transcriptional activity and that CEACAM1 isoforms differ in their inhibitory signaling properties (Maratheftis et al., 2006; Dery et al., 2024). These examples suggest that changes in transcript composition could affect immune and epithelial regulation without substantial changes in total gene expression. If similar alterations occur among other DS-only genes, they may represent an additional source of molecular variation contributing to IBD subtypes that conventional expression analyses would miss. However, our investigations also highlighted the limitations of junction-level evidence, as the observed changes could not be explained by simple exon skipping, and corresponding changes in full-length transcript abundance could not be established. These findings underscore the importance of resolving transcript structures and their potential functional consequences rather than relying solely on statistical significance or junction-level effect size. Extending such analyses to other DS-only genes, particularly those involved in epithelial adhesion, cytoskeletal organization and secretory processes identified in the direct CD vs. UC comparison, may reveal additional mechanisms contributing to shared IBD biology and molecular differences between the two subtypes.

Several limitations should be considered. The greater number of DE and DS genes detected in UC may partly reflect differences in how the two diseases affect the tissue profiled here. UC primarily affects the colonic mucosa, whereas CD is more heterogeneous in anatomical distribution, potentially making disease-associated changes less consistent across colonic samples. Technical differences in sample heterogeneity, sequencing characteristics, residual variation or statistical power that are not related to actual biological differences could also contribute. Thus the increased difference in the UC samples should not be over-interpreted. The limited overlap between DE and DS genes depends in part on their respective significance and effect-size thresholds, while enrichment analyses of DS genes may be affected by differences in the number of testable splicing features per gene (Yan et al., 2018). More generally, bulk RNA-seq cannot distinguish cell-intrinsic regulation from differences in cellular composition, and LeafCutter measures relative intron usage rather than full-length transcript abundance (Y. I. Li et al., 2017). Because intron clusters are defined and filtered separately for each comparison, the absence of a DS call does not necessarily establish subtype specificity, as illustrated by *IRF1*. Although long-read sequencing supported the presence of candidate transcript structures for *IRF1* and *CEACAM1*, only six deeply sequenced biopsies were examined for these genes, and the limited sample size and differences in inflammation status precluded reliable assessment of disease-associated isoform abundance.

Future studies integrating larger-scale long-read isoform sequencing with single-cell transcriptomics will be important for identifying the transcript structures and cellular origins underlying the splicing changes detected here. Larger long-read datasets could more directly test disease-associated differences in full-length isoform usage, while functional validation will be needed to determine whether these changes alter protein output or activity, including the potential effects of alternative transcript initiation on *IRF1* function and exon 7 exclusion on *CEACAM1* signaling. Extending these approaches to the broader set of DS-only genes, particularly those involved in epithelial structural and secretory processes, may help establish whether altered transcript processing contributes to shared IBD mechanisms or subtype-associated differences in epithelial and immune function.

## Conclusion

This study showed that DE and DS capture complementary transcriptomic features in IBD. The limited overlap between DE and DS genes indicates that many splicing changes are not captured by gene-level expression analysis alone. Detailed analyses of the DS-only genes IRF1 and CEACAM1 illustrated the potential functional relevance of altered splice-site usage and transcript structures in IBD. These findings support an integrated DE and DS analysis as a comprehensive framework for identifying disease-associated molecular mechanisms.

## Methods

### Patient recruitment and sample collection

This study received approval from the Human Research Ethics Committee at UNC-Chapel Hill and was conducted in compliance with protocols 10-0355 and 11-0359. Patients were recruited from the UNC Multidisciplinary Center for IBD, with all participants providing written informed consent. Approximately 53% of the participants were female, and the mean age at the time of sample collection was 45 ± 15.6 years. Colonic mucosal samples were obtained from macroscopically uninvolved regions of the ascending colon in IBD patients undergoing colonoscopy or surgical resection. To prevent RNA degradation, tissue samples were immediately preserved in RNAlater and snap-frozen in liquid nitrogen. After sample processing and quality control, a total of 71 non-IBD samples and 225 IBD patient samples, including 126 CD patients and 99 UC patients, were selected for this study.

### Splicing Junction and Gene Expression Quantification

Total RNA was extracted primarily from colonic mucosa samples for paired-end RNA sequencing. Reads were aligned to the human genome (hg38/GRCh38) using GENCODE v47 annotations and processed with STAR (v2.7.11b). Gene-level expression was quantified using HTSeq-count following STAR alignment for each RNA-seq sample. Gene identifiers were harmonized across all samples, and the resulting counts were merged into a unified gene-by-sample matrix containing 78,724 genes quantified across 296 samples.

Alternative splicing quantification was performed using LeafCutter (v0.2.9) (Y. I. Li et al., 2017), which detects differential intron excision events independent of transcript annotation. Splice junctions were first extracted directly from STAR-aligned BAM files using regtools (v0.0.1) junctions extract with default LeafCutter-recommended parameters requiring an 8-bp exon overhang and an intron length between 50 bp and 500 kb. Intron junctions were merged across samples using LeafCutter into intron clusters, grouping junctions that shared common genomic start and end coordinates. Clustering required a minimum of 50 split-read supports per cluster based on LeafCutter-recommended parameters. For each cluster, LeafCutter then computed intron excision ratios, which quantified the proportion of reads supporting each intron relative to all introns in the cluster, forming the percent-spliced-in (PSI) metric.

PSI matrix generation was performed separately for each disease comparison: The CD vs non-IBD comparison group included 93,931 intron junctions grouped into 28,152 intron clusters; the UC vs non-IBD comparison group included 66,600 junctions grouped into 20,803 clusters.

Quality control (QC) of the gene-level expression and splicing matrices was performed using eQTLQC (Wang et al., 2021) to detect outliers across the 296 samples. For expression data, raw count matrices were normalized using the trimmed mean of M-values in edgeR and transformed to log-CPM. For splicing data, LeafCutter intron excision ratios were used directly. The 10,000 most variable introns were retained and no further transformation was applied. Outlier detection was carried out using three complementary approaches: (i) Relative Log Expression analysis, flagging samples with extreme interquartile ranges; (ii) hierarchical clustering of top-variance features coupled with Mahalanobis distance, identifying samples with significantly abnormal profiles; and (iii) the D-statistic, defined as the median correlation of each sample with all others, with the lowest-correlating fraction flagged as outliers. Only samples consistently identified across these methods were excluded. After QC, all 71 non-IBD samples and 225 IBD patient samples (126 CD samples and 99 UC samples) were retained for downstream analyses.

### Batch correction with RUV-seq

To account for technical variation arising from RNA-seq library preparation and sequencing, we used Remove Unwanted Variation implemented in RUVSeq (v1.42.0, R/Bioconductor). Latent factors capturing unwanted variation were estimated with the RUV function separately for the gene expression and percent spliced in (PSI) matrices. For gene expression, raw counts were first variance-stabilized using DESeq2 (v1.48.1, R/Bioconductor), and genes were ranked by across-sample variance; the 500 least variable genes were used as negative control features for the RUV function. For splicing, RUV was run on log-CPM-transformed junction counts after removal of sex and disease effects, with the 10% of junctions with the lowest variance used as control features. In each case, we estimated up to 10 latent factors representing unwanted variation.

To determine the number of latent factors to retain in downstream analyses, we evaluated models with increasing numbers of RUV factors. For differential expression, DESeq2 models included sex, disease status and 0–10 estimated RUV factors as covariates. The optimal number of factors was defined as the point beyond which additional factors did not increase the number of significantly differentially expressed genes, yielding five factors for both the CD vs non-IBD and UC vs non-IBD comparisons. For differential splicing, LeafCutter models were fitted analogously with sex, disease status and 0–10 estimated RUV factors, and the final number of factors was selected to maximize the number of significantly differentially spliced clusters, yielding three factors for the CD vs non-IBD comparison and two factors for the UC vs non-IBD comparison.

### Differential expression analysis

Gene-level differential expression was performed using DESeq2 (Love, Huber, and Anders, 2014). Genes with at least five counts in fewer than 25% of samples were excluded. Analyses were conducted on a count matrix that retained chrX but excluded chrY. Differential expression was assessed using the likelihood ratio test, which compares a full model that includes disease status with a reduced model that excludes it. This approach identifies genes whose expression levels are significantly explained by disease status, after accounting for sex and RUV-derived covariates. We evaluated two primary contrasts, UC vs. non-IBD and CD vs. non-IBD. Genes were considered differentially expressed if they showed an absolute log_2_ fold change greater than 1 and an adjusted p-value less than 0.05. Gene symbols were annotated according to GENCODE v47.

For visualization of global expression structure, PCA was performed on the variance-stabilized expression matrix generated with DESeq2. Samples were plotted according to the first two principal components and colored by disease status. Volcano plots were generated from the DESeq2 results, with log_2_ fold change on the x-axis and the negative log10-transformed adjusted p-value on the y-axis. Genes were highlighted according to significance and effect-size thresholds.

### Differential splicing analysis

Differential alternative splicing was analyzed using LeafCutter, which identifies differential intron excision events based on the clustering of splicing junction counts. Similar to differential expression analysis, the estimated RUV factors were incorporated as covariates for LeafCutter input. Because LeafCutter can only test pairwise contrasts, analyses were performed separately for each disease comparison of interest: CD vs non-IBD and UC vs non-IBD using the corresponding raw count matrix and covariate file as input. Annotation of introns and splice sites was based on GENCODE v47 gene annotation. For this study, differentially spliced clusters were defined as intron clusters mapping to a single gene with a cluster-adjusted *p*-value below 0.05 and a maximum *|*ΔPSI*|* meeting or exceeding the 90th percentile of *|*ΔPSI*|* across all tested introns in the corresponding comparison. The resulting effect-size thresholds were 0.0291 for CD vs. non-IBD and 0.0493 for UC vs. non-IBD. For each qualifying cluster, the intron with the largest absolute ΔPSI was used to represent the splicing event.

For visualization of global splicing structure, PCA was performed on a normalized junction-count matrix generated from raw intron junction counts by library-size normalization in edgeR (v4.6.2) followed by log-CPM transformation. Samples were plotted according to the first two principal components and colored by disease status.

### Long-read sequencing and transcript quantification

Six colonic mucosal biopsies were subjected to long-read RNA sequencing on the Oxford Nanopore platform. Reads were aligned to the GRCh38 primary assembly using minimap2 (v2.31) (-ax splice -k14 --cs=long) in a two-pass workflow. For each sample, splice junctions identified in the first-pass alignments were scored and filtered using 2passtools score and 2passtools filter, respectively. The retained junctions were supplied to the second pass alignment using the --junc-bed option. Alignments were converted to BAM format, coordinate-sorted, and indexed using samtools. Second pass BAM files from all samples were analyzed together using bambu (v3.13.1) to estimate transcript-level read counts. This analysis used the GRCh38 reference genome and GENCODE v50 transcript annotation. The annotation was imported using GenomicFeatures and prepared using bambu::prepareAnnotations, with otherwise default bambu parameters.

### Transcript and junction visualization

For selected genes, annotated transcript structures were visualized using ggtranscript and ggplot2. The exon coordinates and transcript identifiers were obtained from the GENCODE v50 GTF annotation. Pfam protein domain annotations were downloaded as a BED12 file from UCSC genome browser and displayed as individual genomic blocks alongside the transcript structure.

### Pathway Enrichment Analysis

Pathway enrichment analysis was performed separately for DE and DS gene sets using Enrichr through the enrichR package (v3.4) with GO Biological Process 2026, GO Cellular Component 2026, GO Molecular Function 2026 and Reactome Pathways 2024. For DE enrichment, the background comprised 18,434 expressed genes retained after pre-DESeq2 filtering (at least 5 counts in more than 25% of samples) and with an assigned gene symbol, with the same background used across contrasts from the single DESeq2 model. For each library, the background was restricted to annotated genes. Enrichment was assessed by Fisher’s exact test with Benjamini–Hochberg correction within each library, and pathways with adjusted *p <* 0.05 were considered significant.

Significant pathways from the CD vs. non-IBD and UC vs. non-IBD comparisons were further stratified into nine groups according to disease comparison and molecular evidence type. Disease categories were defined as CD-only, UC-only, or shared, and molecular evidence categories were defined as pathways enriched among genes upregulated in disease (DE-up), genes downregulated in disease (DE-down), or DS genes. Within each subgroup and gene-set library, pathway redundancy was reduced using simplifyEnrichment (v2.6.0). For GO terms, pairwise semantic similarity was calculated using the Rel measure implemented in simona (v1.10.0) with annotations from org.Hs.eg.db (v3.23.1). Reactome pathways were compared using Jaccard similarity based on overlap between their annotated gene sets. Related pathways were grouped using the binary cut algorithm, and the pathway with the smallest adjusted *p*-value within each cluster was selected as the representative term. This procedure yielded 141 representative pathways.

To visualize relationships among enriched pathways, EnrichmentMap-style networks were constructed in R using igraph (v2.3.3). For each disease comparison, significant pathways from all four libraries and from DE and DS analyses were combined into a single network using the complete gene-set definitions from the corresponding Gene Matrix Transposed libraries. Nodes represented enriched pathways, and edges connected pathway pairs with a Jaccard similarity of at least 0.35 based on overlap between their annotated gene sets. Network modules were identified using the Markov Cluster Algorithm (MCL v1.0) with an inflation parameter of 1.8. Modules containing at least two pathways were outlined in the network figures, whereas unclustered pathways were displayed as individual nodes. Modules containing at least three pathways were manually assigned biologically representative labels. As part of the module-level comparison, a labeled module was considered shared between the CD and UC networks if its overlap coefficient with any labeled module in the other network was at least 0.5, where the overlap coefficient was defined as the number of shared pathways divided by the size of the smaller module. Shared modules were indicated in green. Network layouts were generated using the Fruchterman–Reingold algorithm and rendered with ggplot2 and ggforce.

## Supplementary Information

### Supplementary Figures

**Supplementary Fig. 1.**
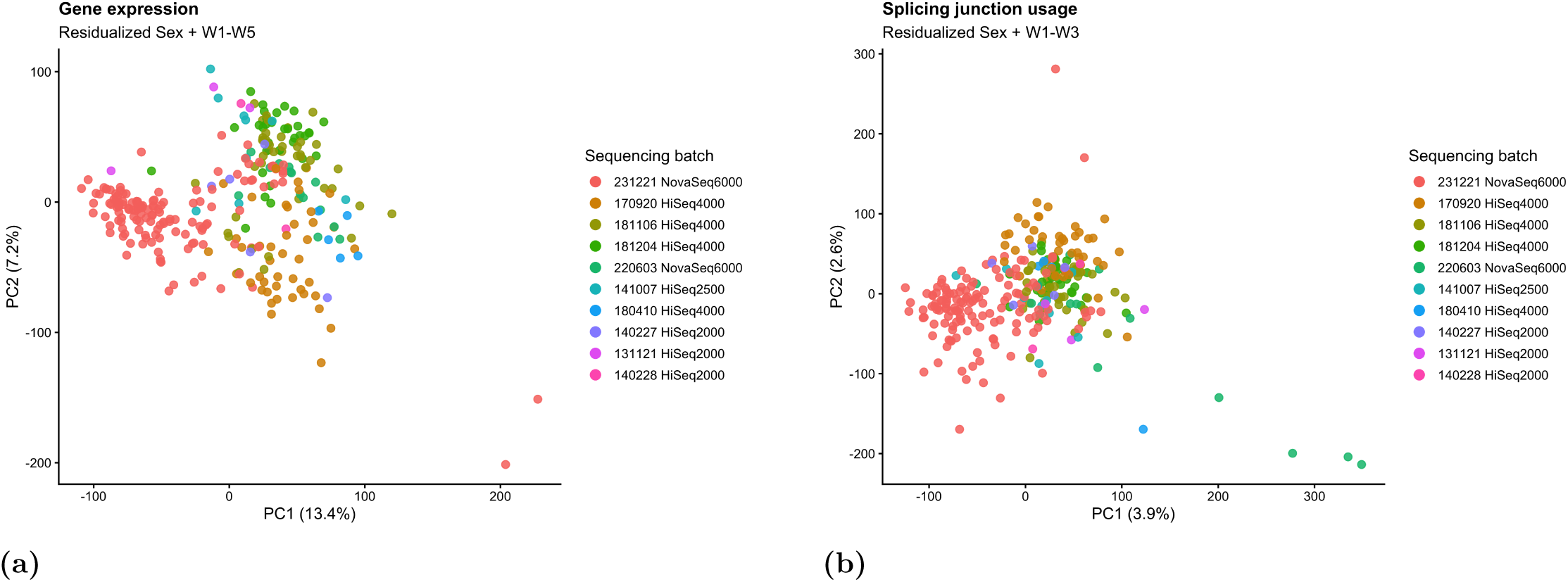
Batch assessment for expression and splicing analyses. **a)** Principal component analysis of the residualized, normalized gene-expression matrix colored by batch-related variables. Sex and the first five unwanted-variation factors (W1–W5), estimated using RUVSeq, were regressed out before visualization. **b)** Principal component analysis of the residualized, normalized junction-count matrix colored by batch-related variables. Sex and the first three unwanted-variation factors (W1–W3), estimated using RUVSeq, were regressed out before visualization. Each point represents one sample.

